# Prediction of drug-protein interaction and drug repositioning using machine learning model

**DOI:** 10.1101/2020.07.29.218826

**Authors:** Yu-Ting Lin, Sheh-Yi Sheu, Chen-Ching Lin

## Abstract

**Background:** Traditional drug development is time-consuming and expensive, while computer-aided drug repositioning can improve efficiency and productivity. In this study, we proposed a machine learning pipeline to predict the binding interaction between proteins and marketed or studied drugs. We then extended the predicted interactions to construct a protein network that could be applied to discover the potentially shared drugs between proteins and thus predict drug repositioning.

**Methods:** Binding information between proteins and drugs from the Binding Database and the physicochemical properties of drugs from the ChEMBL database were used to build the machine learning models, i.e. support vector regression. We further measured proportionalities between proteins by the predicted binding affinity and introduced edge betweenness centrality to construct a protein similarity network for drug repositioning.

**Results:** As the proof of concept, we demonstrated our machine learning approach is capable of reflecting the binding strength between drugs and the target protein. When comparing coefficients of protein models, we found proteins SYUA and TAU that may share common ligand which were not in our training data. Using the edge betweenness centrality network based on the prediction proportionality of protein models, we found a potential target, AK1C2, of aspirin and of which the binding interaction had been validated.

**Conclusions:** Our study could not only be applied to drug repositioning by comparing protein models or searching the protein-protein network, but also to predict the binding strength once the sufficient experimental data was provided to train the protein models.

## Background

In pharmacology, drugs can be classified into several classes by their sizes, mechanisms of action, chemical structures or other different criteria. Among these classes, a small molecule drugs (SMD) often refers to an organic compound with low molecular weight (< 900 dalton) and small size. Compared to biological drugs, SMDs have simple and well-defined structures, leading to easier synthesis and a more predictable chemical process in manufacturing and characterizing. In general, the development of one SMD is mainly processed via 5 stages: I. *Discovery and Development,* II. *Preclinical Research*, III. *Clinical Research*, IV. *FDA Review*, and V. *FDA Post-Market Safety Monitoring*, to make sure that the new pharmaceutical compounds are safe to be distributed to the market [1, 2]. Among the cycle of drug development, the first step, drug discovery, is one of those that cost the most, up to 674 US$ millions per one approved drug and taking 2~5 years in average [3–5]. To boost the process, novel techniques were proposed in recent years, like combinatorial chemistry methods for compound formulation as well as using virtual screening and high-throughput screening to filter out candidate drugs. By using a computational, data-based, or robotic method, the accuracy and efficiency of drug discovery were highly improved. Also, drug repositioning plays an important role in accelerating drug development as well.

Drug repositioning (a.k.a. drug repurposing) has increasingly become an essential strategy in drug development by reinvestigating existing de-risked and approved drugs for new therapeutic purposes [6]. Despite some remaining technological and regulatory challenges needed to be solved, this novel strategy is more efficient, economical, and riskless. On the other hand, with a great amount of biochemical, pharmacological, and clinic data collected in these decades, there have been many emerging techniques like machine learning and network analysis that can take advantage of big data and promote efficiency and productivity [7].

So far, studies using machine learning approaches for drug repositioning have become more and more popular [8–10]. Napolitano F, et al. (*J Cheminform*, 5, 30, 2013) proposed a method based on an SVM (support vector machine) classifier to predict the corresponding disease classes of drugs [11]. Kim E, et al. (*BMC Bioinformatics*, 20, 247, 2019) constructed models using multiple classification algorithms, including logistic regression, random forest, and SVM to predict potential indications for existing drugs [12]. The method developed by Zhanchao Li (*Anal. Methods*, 10, 4152-4161, 2018) uses the random forest algorithm to predict the binding affinity between a compound and a protein directly. Based on the graph theory, the model uses compound-compound similarity, protein-protein interactions, and compound-protein interaction to characterize the binding interaction and may be applied to high-throughput virtual screening [13].

In this study, we employed the support vector regression (SVR) algorithm to build models to predict the binding interaction between proteins and drugs. The inhibition constant (K_i_), which represented the binding strength, was used as label; and structural fingerprint format and physicochemical properties of drugs were used as features. Consequently, we selected 556 protein models to forecast the binding strength with 86,475 drugs. Additionally, we adapt the similarity of predicted binding strength to drugs between proteins to construct a protein similarity network. Furthermore, we proposed an auxiliary network constructed from edge betweenness centralities (eBC) to discover the potential drug repositioning between proteins. With the aid of the network, we found a potential target protein AK1C2 of aspirin by its known target PGH1 and the binding interaction had been reported, providing a successful application of our method for drug repositioning. Briefly, our method can not only be applied to drug repositioning by comparing protein models or searching the protein-protein network, but also by predicting the binding strength directly once the experimental data are sufficient to train the protein models.

## Methods

### Information of proteins and drugs

To predict the strengths of unknown binding interactions of proteins and drugs, we needed experimental data of known protein-drug interactions train the machine learning model. In this study, the protein-drug interaction data were downloaded from the Binding Database [14–18] and interactions with non-human proteins were discarded. Next, the drug compounds are provided with the structural information, PubChem CID, InChIKey, and the binding strength to the target protein, which is inhibition constant in this study, were recognized as informative and kept. For proteins, only human proteins with specific UniProt entry names were adopted. Among the information, the binding strength is the label that our machine learning model attempts to predict; and structural information is one part of the feature in the model. Noted that in our study, only drug information was used as features to describe the binding interactions in each protein model. The circumstances of multiple binding sites of a protein were not considered and only the data with the highest value of binding strength was kept if there were more than one data of a drug-protein pair.

The structures of drugs were described as “Molecular ACCess System (MACCS) keys” [19–21] that consists of a list of binary values (0 or 1) and contains 166 key descriptors to encode a structure. The MACCS keys of drugs were used as a 166-bit feature to train the machine learning models.

On the other hand, since the property data describing physicochemical characteristics of a drug could affect the binding between drug and target protein, we further add them as features in our model. The physicochemical properties of drug compounds were obtained from the ChEMBL database. We used the Python library “chembl_webresource_client”, which is officially developed and supported by the ChEMBL group and selected 17 properties that might have relevance with binding and thus help in prediction. The properties are introduced following their names used in the library (Table S1). Finally, the 17 physicochemical properties were combined with the 166-bit structural fingerprint to form 183 features that were used in our machine learning model.

### Binding strength of drug-protein interactions

In this study, we used −logK_i_ as label to represent the binding strength of a drug-protein interaction and to train the protein models. In fact, the Gibbs Free Energy (ΔG^0^) is also frequently used to represent the binding affinity of a drug and target. From the aspect of thermodynamics, the lower the ΔG^0^ is, the stronger the interaction is. And ΔG^0^ can be calculated by an equilibrium constant (*K*_*eq*_) as Δ*G*^0^ = −*RT*ln*K*_*eq*_, where *R* is the ideal gas constant that equals 8.314 J mol^−1^ K^−1^ and T is the temperature on the Kelvin scale [22]. The equilibrium constant is the dissociation constant in this case, then the ΔG^0^ can be further represented by *K*_*i*_ as 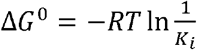. Because *R* and *T* are both constants in the equilibrium, we can infer that ΔG^0^ is positively proportional to log*K*_*i*_ by multiplying ln*K*_*i*_ by a constant log*e*. For the reasons mentioned above, we chose log*K*_*i*_ to represent the strength of the binding of a drug-protein interaction.

### Construction of protein models

In this study, we aim to develop a machine learning model to predict the binding affinity of a drug to a specific protein. The machine learning algorithm we chose here is support vector regression (SVR), which is a regression version of support vector machine (SVM) and works by finding a hyperplane that can minimize the distance from the hyperplane to the instance and cover as most data points as possible in the same time [23]. Because the model of SVM or SVR can be recognized as a weighted sum of the support vectors, it has advantages of reduced computational time and storage requirements over other frameworks. And the principle gives SVM or SVR good generalization capability and better robustness over those algorithms that involve randomness elements.

To train our prediction model, we obtained 152,112 binding information among 1,042 proteins and 86,475 drugs from the BindingDB (Dataset S1) [14–18]. The drug in each data was described by 183 features which consisted of a 166-bit MACCS structure fingerprint [19–21] and 17 physicochemical properties (Table S1) from the ChEMBL database (Dataset S2) [24, 25]. Here we introduced inhibition constant (*K*_*i*_) to represent the binding strength of a drug with its target protein. *K*_*i*_ is defined as 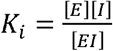 where [E], [I], and [EI] are the equilibrium concentrations of enzyme, inhibitor, and the complex [26]. The smaller the *K*_*i*_ value, the greater the binding strength of the inhibitor with its target, and the better the potency of the drug in most cases. Since the *K*_*i*_ value can quantitatively reflect how tightly a compound binds to a particular protein, we used it to represent the binding strength of a compound-protein interaction. In our data, however, the experimental *K*_*i*_ values show an exponential distribution and many of the experimental *K*_*i*_ values are larger than 1,000 while some of the others are less than one. Given that many regression algorithms are based on the concept of least squares method to approximate data, the large difference in scale between observations leads to poor fitting since the residues on the larger scale weigh moreover those on a smaller scale. Therefore, we transformed the experimental *K*_*i*_ values into logarithmic ones, to offer the machine learning model a dataset with a linear distribution. Furthermore, to make the study results easier to be comprehended, we used −logK_i_ as the indicator of binding affinity and the label of our machine learning model, while the larger the value is, the stronger the compound-protein interaction is.

Before model training, each property feature among samples were standardized by computing the standard score z as *z* = (*x* − *μ*)/*σ* where *μ* is the mean and *σ* is the standard deviation of the feature throughout all samples. We then built a prediction model for each protein using only the dataset targeting it. The model trainings were all default settings except the parameter “kernel” was specified to be “linear”. When training the SVR model, we randomly split the dataset into two groups: 70% of the dataset as the training set to build the model and evaluate the prediction accuracy, and the other 30% as the testing set for validation to assess how the model would generalize to an independent dataset. Then the models applied for further analysis would be trained by their complete datasets.

The evaluation we used here to see how well a model could predict was the Spearman correlation coefficient (SCC, *ρ*), which is a nonparametric measure of rank correlation to assess the monotonic relationships [27–29]. When comparing the accuracies among models, we introduced the Fisher transformation to take the data number into account [30]. The transformation of the sample SCC value *r* can be computed by

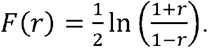

When given the sample size *n*, the *z*-score of *r* is

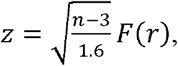

which approximately follows a standard normal distribution under the null hypothesis of statistical independence (*ρ* = 0).

### Construction of the protein network for drug repositioning

Having the predictions of binding strength, the protein similarity was calculated by proportionality [31–33]. However, since the protein similarity only considers the direct association between them, we further constructed an auxiliary network which was based on proportionalities and was capable of detecting the indirect similarity between proteins.

To keep the highly confident protein pairs, we first removed the protein pairs with low proportionality, which is the majority of the network, and applied the concept of the shortest paths in the network to consider the indirect correlations since the two adjacency proteins might not have the strongest correlation when computing direct similarity.

The concept of the shortest path was applied in our study by betweenness centrality, which is used to evaluate the importance of nodes of edges in a network. If a network is weighted, the shortest path is the one having the maximal sum of the weights of the edges. The edge betweenness centrality (eBC) of an edge is defined by the number of these shortest paths that pass through this edge and the eBC of an edge *v* can be computed as

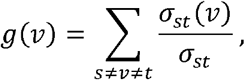

where σ_st_ is the sum of the shortest path from node s to node t, and *σ*_*st*_(*v*) is the number of the paths that pass through *v* [34].

For our original network of proportionality, we calculated the eBC of every edge in it and removed those whose eBC was 0. Then each remained edge was assigned a new weight by its eBC value for next eBC calculation. The cycle of eBC calculation, removing edges, and assignment of new weights was repeated until convergence (Figure S1A). Here we used the mean-square error (MSE) between the edge weights and the eBC values in a cycle to tell whether the calculation was converged. The number of remained edges maintained the same after the 15^th^ cycle and so did the MSE after the 18^th^ (Figure S1B). We then used the eBC values calculated in the 18^th^ cycle to build an eBC-based protein-protein network for further analysis.

## Results

### Proof of concept for the machine learning model

To determine the capability of our machine learning pipeline in predicting the binding affinity of a drug to a specific protein, we first used the models trained from AA2AR, DHI1, and AL5AP separately as a proof of concept. First, we partitioned our dataset into training and testing sets to perform cross-validation for these three proteins. We observed that the experimental −*logK*_*i*_ values were strongly and positively correlated with the predicted −*logK*_*i*_ values from training and testing sets (Table 1 and Figure 1a). These results implicated that this approach might be capable of reflecting the binding strength of a drug to target protein well to some extent. However, the process of random sampling for cross-validation makes deciding the best model uncertain. Additionally, to get the best performance when given an unknown drug, the complete dataset targeting a specific protein should be used to train the model. Hence the accuracy and the risk of overfitting using the complete dataset to train and test the model were examined (Figure 1a). The SCC values of complete datasets in the three models were close to the distributions of the training sets as well as testing sets (Table 1), demonstrating the low risk of overfitting from complete dataset. Though the SCC values themselves may be affected by the number of drugs, leading to lower accuracies of the complete datasets than the training sets, the models built by complete datasets were still representative for predictions. Thus, the models in the following study were trained by their own complete dataset for best predictions.

**Table 1.**
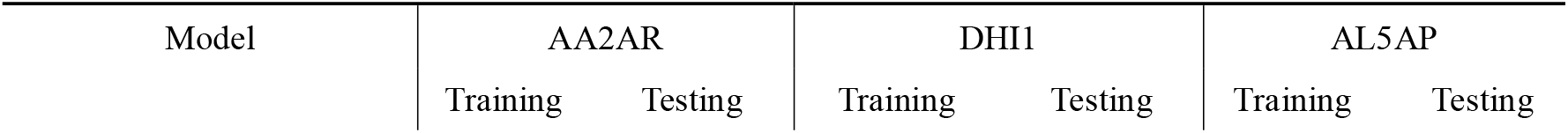
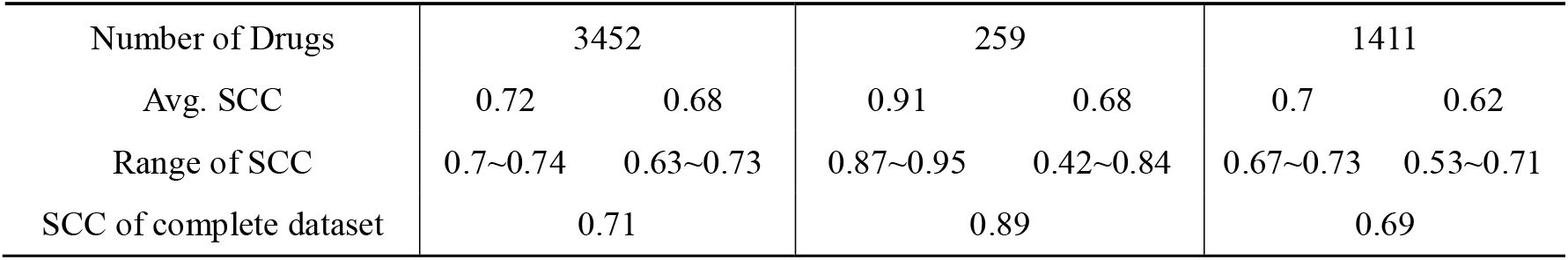
Performance of three models: AA2AR, DHI1, and AL5AP.

| Model | AA2AR |  | DHI1 |  | AL5AP |  |
| --- | --- | --- | --- | --- | --- | --- |
|  | Training | Testing | Training | Testing | Training | Testing |
| Number of Drugs | 3452 |  | 259 |  | 1411 |  |
| Avg. SCC | 0.72 | 0.68 | 0.91 | 0.68 | 0.7 | 0.62 |
| Range of SCC | 0.7~0.74 | 0.63~0.73 | 0.87~0.95 | 0.42~0.84 | 0.67~0.73 | 0.53~0.71 |
| SCC of complete dataset | 0.71 |  | 0.89 |  | 0.69 |  |

**Figure 1.**
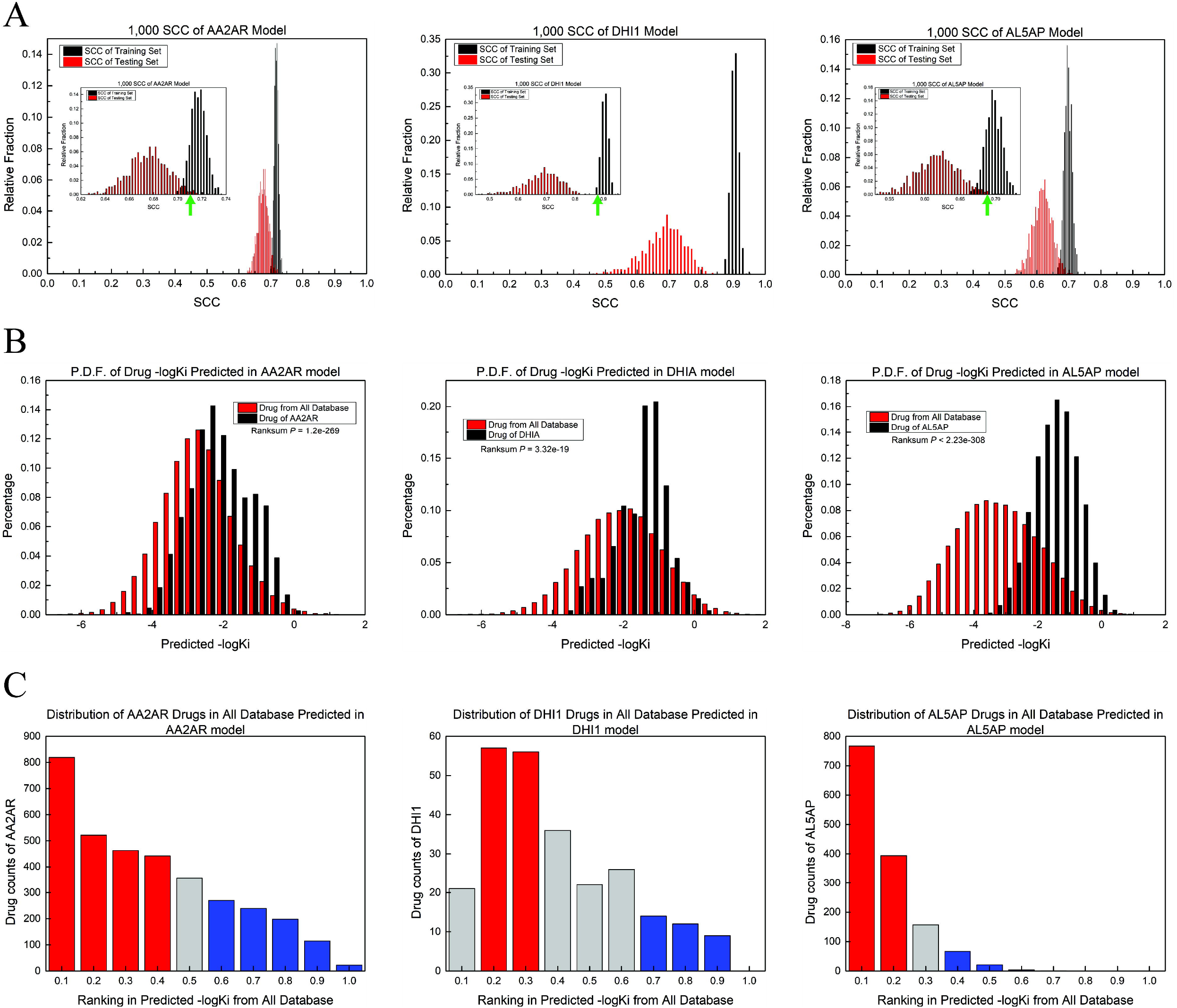
Performance of the prediction model from AA2AR, DHI1, and AL5AP protein. a. The distribution of SCCs between experimental and predicted binding affinity from trained (black) and tested (red) models of AA2AR, DHI1, and AL5AP. The green arrow indicated the SCC of the complete dataset. b. Distributions of experimental (black) and predicted (red) binding affinity of drugs to AA2AR, DHI1, and AL5AP. c. Number of experimentally-validated drugs of AA2AR, DHI1, and AL5AP in ascending rankings of predicted binding affinity. The red and blue bars showed the experimentally-validated drugs in the corresponding ranking are significantly over- and under-represented, respectively (Fisher’s exact test, *p*-value < 0.05).

We further examined the ability of a protein model to select the drugs that would have binding interaction with the corresponding protein from all the drugs in the dataset. For example, the −*logK*_*i*_ values of all 86,475 drugs predicted from the AA2AR model were compared with that of AA2AR experimentally-validated drugs. We found that the −*logK*_*i*_ values distribution of experimentally validated drugs were significantly higher than those of other drugs (Figure 1b). Additionally, the experimentally validated drugs were observed to be significantly enriched in the drugs with higher predicted −*logKi*, and the amount decreased over the rankings. (Figure 1b). This supported that our model could not only pick out the drugs that were already known to have binding interaction with the modeled protein but also implicate the unvalidated drugs in the top rankings may have potential binding ability to the modeled protein. To further demonstrate that the models could predict the drugs strongly binding with the modeled protein, we investigated the drugs with high predicted and experimental *logK*_*i*_ to AA2AR. The compound of 3-[2-[4-[2-Fluoro-4-(2-methoxyethoxy)phenyl]piperazin-1-yl]ethyl]-8-(furan-2-yl)-[1, 2,4]triazolo[5,1-f]purin-5-amine is predicted with the secondly highest −*logK*_*i*_ value of 0.094 among the drugs targeting AA2AR and possessed the 99^th^ high experimental −*logK*_*i*_ value of 0.097 in the training set. The experimental *k*_*i*_ value of the compound binding to AA2AR is 0.8 nM, making the compound a promising antagonist to AA2AR. The possible application of the drug could prevent motor disturbances caused by defects related to dopamine D2 receptors and further serve as an anti-symptomatic drug for the treatment of Parkinson’s disease [35]. Briefly, using AA2AR, DHI1, and AL5AP models as exampling proof of concept showed that our method based on the machine learning approach could be used to predict the relative binding strength between a drug and a protein, which could be applied to the screening of candidate drugs in drug development.

### Performance of the machine learning model

After demonstration of the proof of concept for our machine learning pipeline, we then attempted to construct the prediction models for all the proteins in the database. However, the machine learning approaches were known to be affected by the sample size - number of experimentally validated drugs in this study. We also observed that, compared to AA2AR and AL5AP model, DHI1 model, which possessed fewer experimentally validated drugs, showed larger difference of SCC between training and testing set (Figure S2). Therefore, we verified the effect of the training data size on the performances of the prediction models. Among the total 1,042 proteins from the BindingDB, over 46% of them have less than 10 data of binding information with drugs and only 21% with more than 100 data (Figure 2a). Furthermore, we found that as the number of drugs increased, the SCC z-score values of model predictions became higher, suggesting better accuracies (Figure 2b). Also, the more experimentally-validated drug data a protein model was built by, the smaller the standard deviation of the SCC distribution (Figure 2c) and the more unapparent the SCC difference between the model trained by the complete dataset and by the training set (70% of complete dataset) were observed (Figure 2d). These results also indicated that the model trained for the protein with more experimentally validated drugs could be more stable. Accordingly, to prevent models from overfitting and inaccuracies caused by lacking training data, 556 proteins (53% of all) with more than 10 drug data were retained for the following analysis. Among 556 proteins, 513 (92.26%) protein models are provided with SCC z-score greater than two, suggesting the high accuracy of the machine learning pipeline in predicting relative −*logKi* between drugs and proteins.

**Figure 2.**
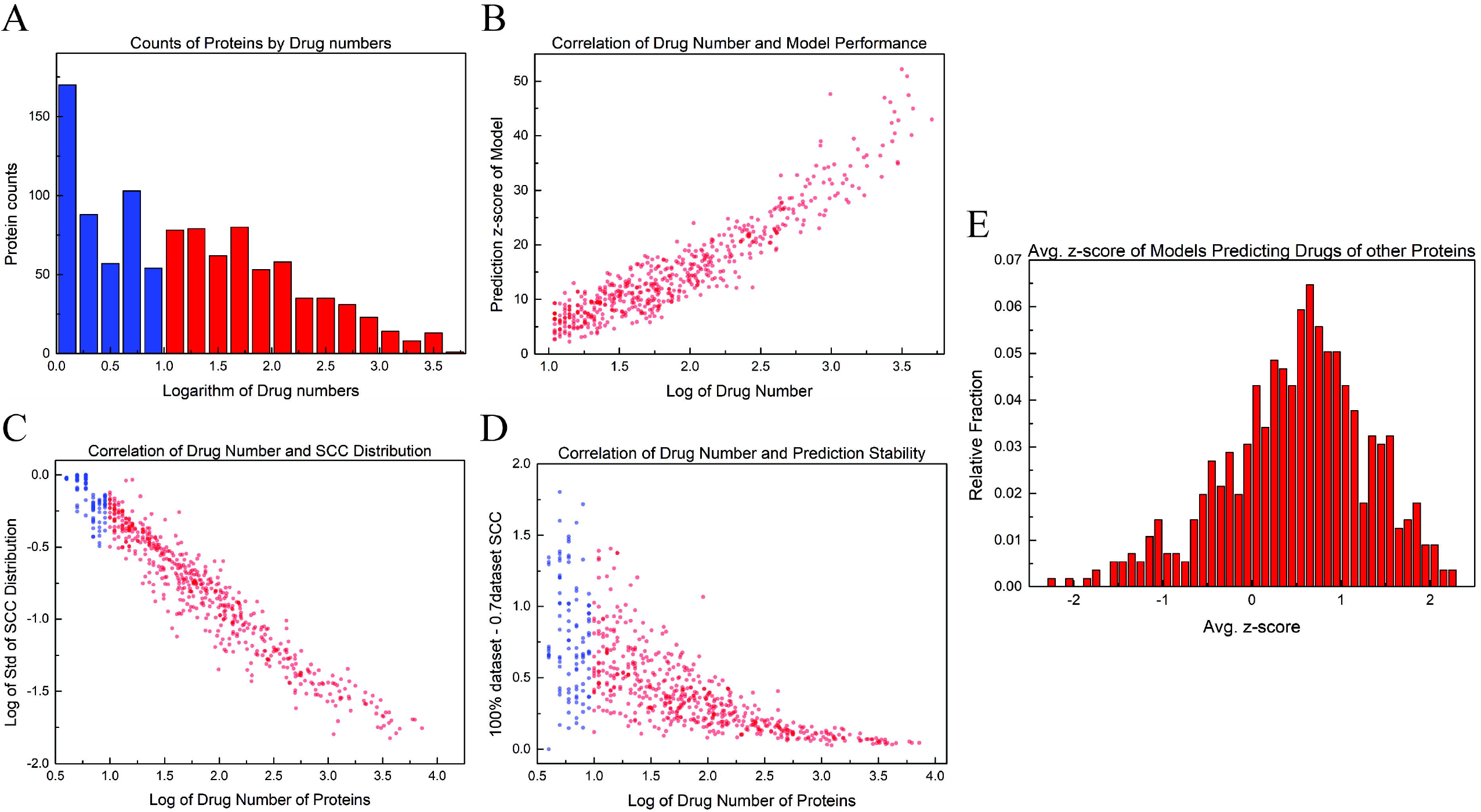
The association between number of experimentally-validated drugs and model performance. a. Distribution of the number of experimentally-validated drugs for proteins. Blue and red bars showed the proteins with experimentally-validated drugs less and more than ten, respectively. b. Positive correlation between the drug numbers and prediction SCC transformed z-score of models. c. Negative correlation between the drug number and standard deviation of SCC. The standard deviation was calculated from the SCCs between the experimentally-validated and predicted binding affinity from 1,000 training model. d. Association between the drug numbers and prediction stability. The prediction stability here is estimated by the difference of SCC between the complete dataset and training set. (E) Average SCC z-score of models predicting drugs of other proteins. Most of the models were unsuitable to predict the binding interaction targeting proteins other than themselves.

Next, we testified the model specificities by using the datasets from the other proteins. For example, to test the AA2AR model, we used it to predict the −*logK*_*i*_ of data originally targeting DHI1 and AL5AP. Then, for DHI1 and AL5AP, we compared their −*logK*_*i*_ predicted by AA2AR model to their experimental −*logK*_*i*_ separately. As expected, the SCC between experimental −*logK*_*i*_ targeting DHI1 and predicted −*logK*_*i*_ from AA2AR model was -0.21; and the SCC between experimental −*logK*_*i*_ targeting AL5AP and predicted −logK_i_ from AA2AR model was 0.09 (Figure S3). The two results suggested that the AA2AR model could not be used to predict the −*logK*_*i*_ of drugs binding to DHI1, neither AL5AP. Same evaluation was performed on DHI1 and AL5AP models and it was alluded that a protein model could only be applied to predict the drugs targeting the protein itself and showing the specificity of the prediction model (Figure S3). The investigation of all the models was further evaluated by the distribution of the average z-score. For all 556 proteins, every model was used to predict the −logK_i_ of other 555 sets of drugs and their target proteins, and the average z-score value was recorded (Figure 2e). Notably, only 1.6% proteins are provided with average z-score greater than two, showing that most of the models were unsuitable to predict the binding interaction targeting proteins other than themselves and assuring the overall specificities of models.

Nevertheless, based on the reason, the specificities of models indicated that if a model could predict the drugs of another protein well, there might be some similarities or shared characteristics between the two proteins that could be applied to drug repositioning. For example, the model of protein DRD2 was trained by 5,155 drug data and had a z-score of 20 when predicting the 844 drugs binding to their experimental target protein BCL2. The two proteins shared significantly enriched GO biological process (p-value < 0.0001, Fisher’s exact test), showing their functional similarity. DRD2 is dopamine D2 receptor whose activity is mediated by G proteins and is the main receptor for most antipsychotic drugs [36]. BCL2 is apoptosis regulator (B-cell lymphoma 2) which can suppress or induce apoptosis and regulate cell death by controlling the mitochondrial membrane permeability [37, 38]. The relevance of the two proteins in several signaling pathways has been studied. In non-small cell lung cancer, it was found that the upregulation of BCL2 is related to the knockdown of DRD2 and the overexpression of the DRD2 gene may decrease the mRNA level of BCL2 [39]. In another research on Parkinson’s disease, it was found that the inhibition of hsa-miR-200a can upregulate BCL2 and DRD2 via the cAMP/PKA signaling pathway and suppress apoptosis in striatal neuron cells [40]. Though no research reported the shared drugs of these proteins, we still demonstrated that using model specificity could help find proteins with similar biological roles and the relevance of them may provide information for medication.

### Model comparison and drug repositioning

We have demonstrated that the SVR models could be able to predict the relative −*logK*_*i*_ value between a drug and a target protein. Therefore, our approach could be a feasible application in compound screening during drug development, especially for repositioning of the approved drugs that are provided with detail and complete experimental information. For example, two proteins with similar models after training might possess similar drug binding characteristics. Accordingly, their models might be able to predict promising drugs as a potential treatment for each other, a.k.a. drug repositioning. We then calculated the SCC of coefficients between models to assess the similarity of drug-binding profiles between proteins. That is, two proteins with higher SCC are more similar to each other on drug-binding characteristics. Accordingly, for each model pair, we inspected the association between the SCC of model coefficients and the average drug number (Figure 3). The results showed that when the models were trained by few drugs, their model similarities varied from 0.89 to -0.56 and when trained by more drugs, the model similarities became more convergent and lied mostly between ±0.2 The *p*-values were examined and the smallest positive SCC value with significance (*p* value < 0.05) was 0.15 (the green line in Figure 3). The proteins of the model pairs above the significance line could be worth to be investigated for drug repositioning. Among all the protein pairs, three of them are with more than 3,000 drugs in average and SCC higher than 0.3. In two of the pairs, CAH1-CAH2 and DRD2-DRD3 were from the same family. Since proteins in a family typically have similar structures, functions, and sequence similarity, it was reasonable for their prediction models with similar feature weight ranking, that is, with high SCC. This also demonstrated that our method could recognize the proteins with some shared characteristics. However, for the proteins in the same family, most of the experimented drugs were the same, limiting the efficacy of repositioning. Thus, we focused on another protein pair, 5HT2A-DRD3. Protein 5HT2A is a G-protein coupled receptor for 5-hydroxytryptamine (serotonin) and can affect neural activity, perception, cognition, and mood. It also plays roles in the regulation of behavior, intestinal smooth muscle contraction, and arterial vasoconstriction [41, 42]. DRD3 is a dopamine receptor whose activity is mediated by G proteins which inhibit adenylyl cyclase and can promote cell proliferation [43]. Among 2,683 drugs of 5HT2A and 3,535 drugs of DRD3, 914 shared drugs are significantly over-represented (p-value < 0.0001, Fisher’s exact test), suggesting that they did share binding interaction that had been under study.

**Figure 3.**
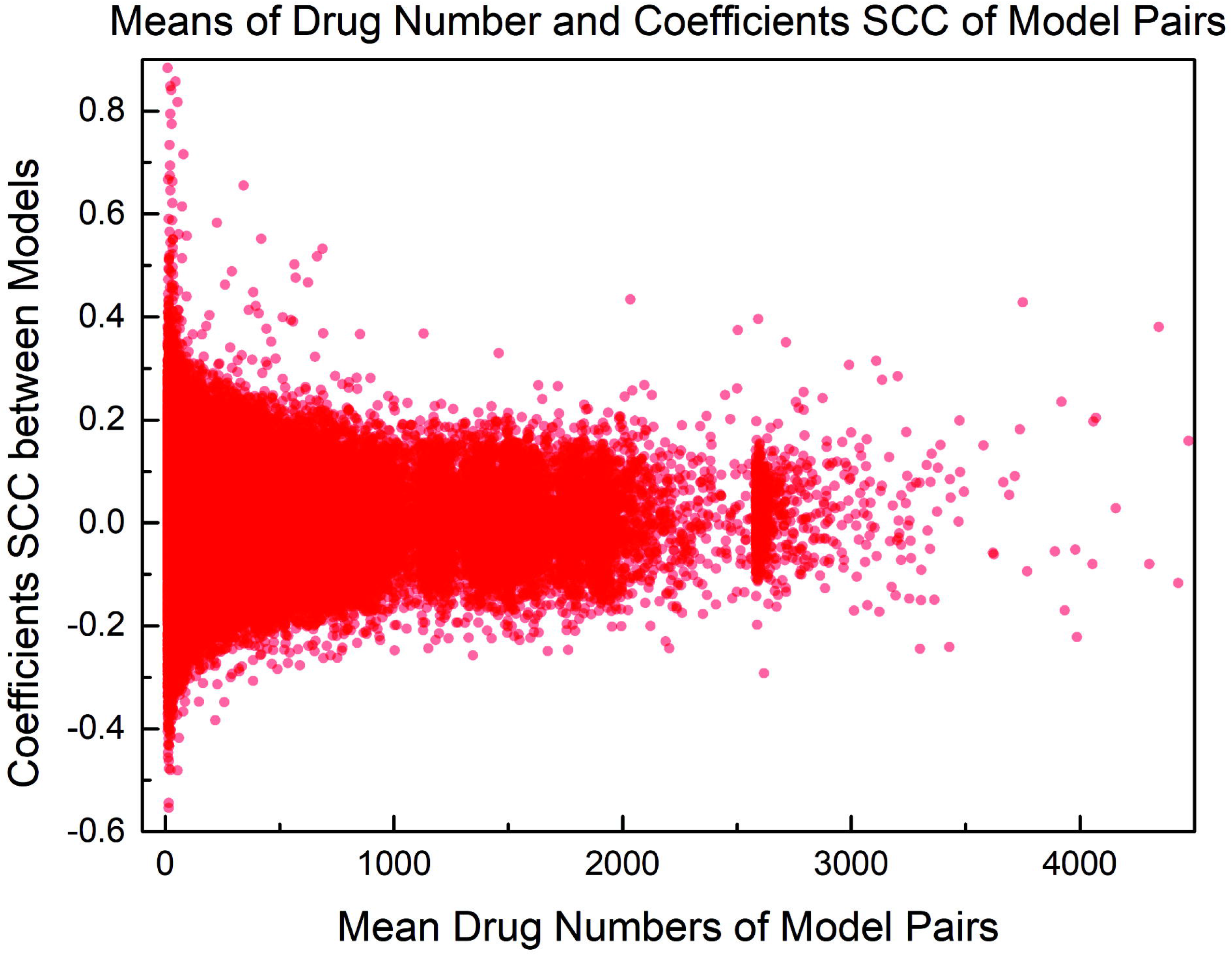
The SCC of model coefficients and the mean drug number of model pairs. For each protein pair, the SCC of model coefficients and average drug number were calculated. The green line indicated where the smallest significantly positive SCC (value ≧ 0.15, *p*-value < 0.05).

However, for the models trained by few drugs, their comparisons may not be convincing enough with a wide range of model coefficients similarities since it was demonstrated that the models performed better with more drugs. Thus, once there were models that were trained by enough data and had high correlations, the two proteins may have alike binding interactions. Another problem was that the difference in the sample size of models may lead to deviations when comparing the coefficients directly, making it inappropriate to build a network by model similarities. Therefore, we proposed the construction of a protein-protein network based on prediction similarity between all protein models to reduce the data amount effect on a single model and provide insights on the mutual connection of proteins.

### Drug repositioning using the eBC network

To apply the protein models to drug repositioning from an integral aspect, we constructed protein similarity networks guided by the SVR model (Figure 4a and 4b). In the network, edges linking proteins are weighted by the proportionality calculated from the predicted −*logKi*. To note, the distances between proteins in the network were assigned by 1 - proportionality. In other words, edges with higher proportionality, i.e. more similar, were assigned by shorter distance. Next, we used the edge betweenness centrality (eBC) to identify the shortest (highest similarity) paths between proteins. Accordingly, the indirect association between proteins, that is, the paths with highest similarity included more than one proteins, could be discovered. Interestingly, the average GO term similarity of protein pairs with a two-step distance was the highest (Figure 4c), meaning that when computing similarities via one intermediate protein allows the eBC network to represent the correlation of the roles that the proteins play in biological processes. Hence it may be feasible for drug repositioning to consider the proteins’ second neighbors in the eBC network if they would have shared drugs and binding interactions.

**Figure 4.**
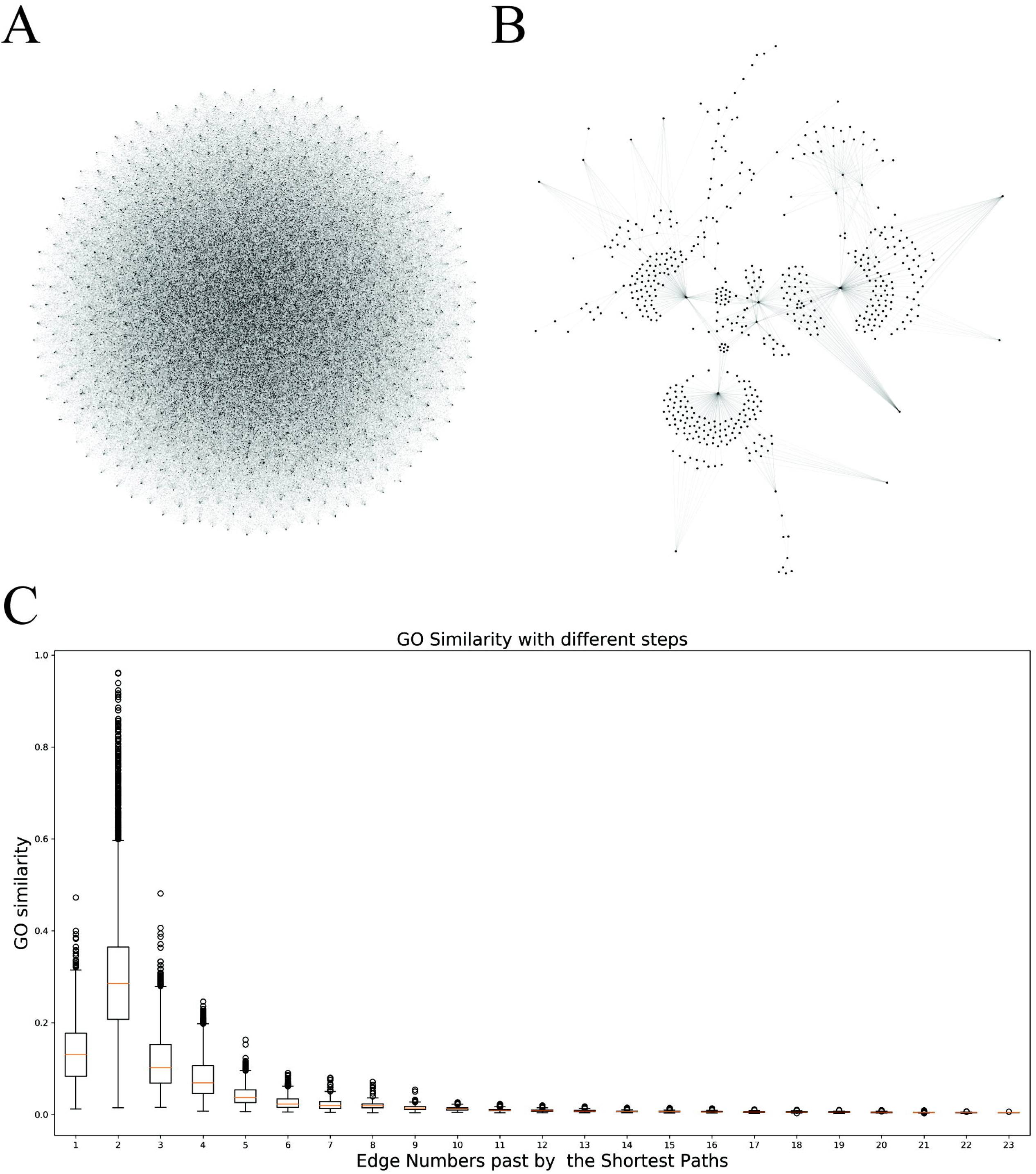
The functional similarity of the shortest paths in the protein similarity network. a. The original network based on protein proportionalities. b. The auxiliary eBC network of the 556 proteins. For the primeval network, the proteins were fully-connected and could not be interpreted with corresponding biological similarities. After removing connections with low edge betweenness centrality, the network had positive correlation with GO term similarities and showed noticeable structure. c. The protein pairs with a two-step distance had the highest GO similarity among pairs with each shortest paths. This implied that the indirect connections between proteins could better reflect the biological similarities.

To determine the reliability of our eBC network for drug repositioning, we chose aspirin (acetylsalicylic acid) as an example to find its potential target protein. Aspirin is a drug widely used to reduce pain, fever, or inflammation, and also applied to prevent strokes and blood clots. One of the known target proteins of aspirin is cyclooxygenase-1 (COX-1), which will be irreversibly inhibited by the acetylation of a serine residue in the active site [44–46]. Basing on COX-1, we then tried to find proteins in the eBC network that may also be targets of aspirin. Among the proteins that were in two-step distance from COX-1, AK1C2 had the highest 2-step GO term similarity with a value of 0.67. AK1C2 is aldo-keto reductase family 1 member C2 that is encoded by gene *AKR1C2*. The enzyme catalyzes the inactivation of 5-alpha-dihydrotestosterone (5-alpha-DHT) to 5-alpha-androstane-3-alpha,17-beta-diol (3-alpha-diol) [47, 48]. The selective loss of AK1C2 may have the relevance of breast and prostate cancers [49, 50] and the overexpression of it may be a high-risk factor in bladder cancer [51]. In our dataset used to train the AK1C2 model, none of the data was about the binding of aspirin to AK1C2, leaving aside the possibility that the network had already had the information of this binding interaction. In a paper studying the inhibition of AK1C2 by nonsteroidal anti-inflammatory drug (NSAID) analogs proposed in 2009 by Michael C. Byrns et al., it was reported that IC_50_ value of inhibition of AK1C2 by aspirin was 16 μM. Also according to the study, the concentrations of aspirin and its deacetylated product salicylic acid needed for inhibition were lower than those of other NSAIDs [52].

The GO term similarity of biological process between COX-1 (PGH1) and AK1C2 in our network was calculated via EST2, a cocaine esterase. Most of their shared GO biological process terms were related to the metabolic process for substances including lipid, organic acid, carboxylic acid, and others. The other terms were mainly related to cellular responses to stimulus. Though without support from known experimental information, it might be possible that there were some correlations between the shared drug and common GO biological process terms of the two proteins, which was worthy of further studying.

## Discussion

We have proposed a machine learning pipeline to discover the potential drug positioning for proteins. To further understand which structure characteristics of drugs affecting the binding with proteins, we analyzed the feature importance of the learning functions. Here we chose three models with the highest SCC *z*-score values of predictions and inspected the absolute values of weights of their physicochemical property features. The protein model with the highest *z*-score of 52.2 among all the models was trained from FA10 by 3,158 binding data. Among 17 physicochemical property features in the FA10 model, the absolute values of their coefficients ranged from 0.05 to 0.89. The feature with the largest absolute weight of -0.89 was the number of non-hydrogen atoms in the molecule, accounted for 10.5% of the 17 features. The one with the second-high absolute value of weight was the polar surface area (0.84, 9.9% of all); the third feature was the molecular weight of the parent form of the molecule (0.83, 9.9% of all). The protein model with the second-high prediction z-score of 50.89 was AA2AR trained by 3,452 data. The absolute values of its features’ coefficients ranged from 0.01 to 1.09. The three top features in descending order of their absolute values of weights were the number of non-hydrogen atoms in the molecule (1.09, 22.3% of all), the count of hydrogens attached to nitrogen or oxygen atoms (0.79, 16.2% of all), and the number of hydrogen bond donors (0.48, 9.9% of all). The third-high z-score of 47.63 belonged to the TRY1 model trained by 986 data. The absolute values of 17 feature coefficients ranged from 0.01 to 1.34. The three top features in descending order of their absolute values of weights were the count of nitrogen and oxygen atoms in the molecule (-1.34, 23.1% of all), the polar surface area (1.01, 17.4% of all), and the count of hydrogens attached to nitrogen or oxygen atoms (-0.518, 8.9% of all). The last model examined was DRD3, which had the forth-high z-score value of 47.43 and was trained by 3,535 data. Its absolute values of 17 feature coefficients ranged from 0.01 to 1.25. The three top features in descending order of their absolute values of weights were the number of non-hydrogen atoms in the molecule (-1.25, 23.4% of all), the value of logP (0.73, 13.8% of all), and the polar surface area (0.69, 13% of all).

Among the four models, three of them had polar surface area (PSA) as an important feature, which is defined as the surface sum over all polar atoms or molecules [53]. In general, bindings between proteins and drugs will be affected by non-bonded interactions, which can be further divided into electrostatic interactions and Van der Waals interactions. While many of the binding interactions depend a lot on the formation of hydrogen bonds and salt bridges, the ligands with more PSA might bring about stronger electrostatic interactions. Also, the PSA property is related to the solvation of a molecule and will affect the inhibition ability of a drug to its target protein. Therefore, PSA played an important role in binding strength prediction and weighed higher in our models [54].

The other features with high weights were also associated with hydrogen atoms and hydrogen bonds, including the number of hydrogen bond donors (HBD), the count of nitrogen and oxygen atoms (HBA Lipinski), count of hydrogens attached to nitrogen or oxygen atoms (HBD Lipinski), and the number of non-hydrogen atoms in the molecule. Unlike PSA, the weights, or coefficients, of these features differ in sign. This is probably because that different target proteins may serve as donors or acceptors of hydrogen bonds, while the number of donors/acceptors of drugs and the binding sites of proteins also contribute to the interaction, leading to diverse prediction pattern in different protein models.

Next, we tried to apply the similarity of feature coefficients between protein models to drug repositioning. Among all 154,290 protein pairs, there were 37 pairs with SCC values of coefficients higher than 0.5 among, and of which each of the 35 pairs was from the same protein family. Since proteins in a family typically have similar structures, functions, and sequence similarity, it was reasonable for their prediction models with similar feature weight ranking, that is, with higher SCC. Also, for the proteins in the same family, most of the experimented drugs were the same, limiting the efficacy of repositioning.

Thus, to further discover the capability of identifying the drug repositioning between proteins from different family, we tried to study the other two protein pairs, SYUA-TAU, and CY24B-WEE1. Since no research of common ligands to these two proteins was found, this makes it difficult to demonstrate the association between CYBB and WEE1. Additionally, the numbers of drug data used to train the models are relatively small, which were 19 and 14 for CY24B and WEE1, respectively. The size of training sets might not be enough for models to approximate precise functions for prediction, and lead to bias when comparing model coefficients. Consequently, we focused on the protein pair, SYUA-TAU. The coefficients of the models of SYUA and TAU had the second-high correlation among all the protein pairs, with an SCC value of 0.86. SYUA is a protein called alpha-synuclein encoded by the *SNCA* gene. It mainly locates in the brain other than other tissues and is critical for brain function [55]. And TAU is a protein named microtubule-associated protein tau, which is encoded by gene *MAPT* [56, 57]. TAU can stabilize microtubules and is abundant in neurons of the central nervous system. Both proteins are associated with pathologies and dementias of the nervous system like Parkinson’s disease and Alzheimer’s disease, in which abnormal aggregations into amyloid of these proteins are often observed. In these disorders, the alpha-synuclein protein and Tau protein form insoluble fibrils that are characterized by Lewy bodies and neurofibrillary tangles respectively [58].

The SYUA model and TAU model were built by 45 and 46 drug data separately, while the 45 drugs were the same and the TAU model had one more drug, FDDNP, in the training dataset. FDDNP (2-(1-{6-[(2-[fluorine-18]fluoroethyl)(methyl)amino]-2-naphthyl}-ethylidene)malono nitril) is a positron emission tomography (PET) molecular imaging probe used to visualize Alzheimer’s disease pathology and its ability to detect amyloid in the brain is already proved. Due to the model similarity between the two proteins, FDDNP might also have binding interaction with SYUA. Additionally, it has been reported that FDDNP has a binding interaction with alpha-synuclein filaments *in vitro* and the *K*_*i*_ value is 210 nM, while in our study the *K*_*i*_ value of FDDNP binding to TAU was 256 nM [59]. The two close experimental *K*_*i*_ values suggested that by comparing model coefficients, we could find the proteins that may have similar binding patterns and even share common ligands. However, FDDNP is majorly used as radiotracers of PET for clinical diagnosis for Alzheimer’s disease and seldom serves as direct medication for treatments. This case demonstrated a promising application in drug repositioning, though it is not the direct usage in treatments.

Our study so far focused on building single model for each protein and used only drug structures and physicochemical properties as features. In the future, an advanced goal is to incorporate features of protein and drug to construct a general model that can predict the binding strength once given a protein-drug pair. Unlike the single models that were restricted to the amount of experimental data of proteins, the general model takes protein features like structures into account and is trained by the binding information targeting all proteins, making it more feasible for prediction of rare protein-drug instances.

## Conclusions

In this study, we applied a machine learning approach to predict the drug-protein interaction and extended the result to construct a protein-protein network for drug repositioning. We demonstrated that the models trained by more drug data had better performance and less risk of overfitting when comparing the SCC values between training and testing results. We further showed that our protein models were able to recognize their own drugs from all the compounds in the dataset, which suggested that the models could be applied for screening of candidate drugs. Also, by comparing model coefficients of physicochemical properties, protein pair SYUA-TAU was found to have alike biological role on GO term annotation and similar binding interactions with shared drugs. When using protein eBC network based on proportionality for drug repositioning, we found a potential target protein AK1C2 of aspirin by investigating the two-step neighbors around its known target PGH1. Briefly, our method can not only be applied to drug repositioning by comparing protein models or searching the protein-protein network, but also by predicting the binding strength directly once the experimental data are sufficient to train the protein models. When given a drug with structure and property information, we can get its binding strengths targeting different proteins and find the one with the highest value as a new medication. Also given a specific protein, we can use its model to predict all the drugs that have experimental data and find those with high returned values for alternative medication. As the accumulation of experimental data and clinical research, we believe that the problem raised by data size will be solved soon in the future and by using our method, the efficiency and cost of drug development can be greatly improved.

## List of abbreviations

SMD: small molecule drugs
FDA: Food and Drug Administration
SVM: support vector machine
SVR: support vector regression
SCC: Spearman correlation coefficient
eBC: edge betweenness centrality
MSE: mean-square error

## Declarations

### • Ethics approval and consent to participate

Not applicable.

### • Consent for publication

Not applicable.

### • Availability of data and materials

The datasets used and analyzed in this study are available in the Binding DB website (https://www.bindingdb.org); the ChEMBL repository (https://www.ebi.ac.uk/chembl); and the Gene Ontology resource (http://geneontology.org/).

### • Authors’ contributions

SYS and CCL directed this study. CCL conceived and designed this study. YTL implemented the computational method and carried out the analysis. YTL drafted the manuscript. SYS and CCL revised the manuscript. All the authors read and approved the final manuscript.

## • Acknowledgements

We thank Dr. Hsuan-Cheng Huang and Dr. Yu-Chao Wang in the Institute of Biomedical Informatics, National Yang-Ming University provided invaluable suggestions in improving this study.

## Reference

1. Ciociola AA, Cohen LB, Kulkarni P, Gastroenterology FD-RMCotACo: How drugs are developed and approved by the FDA: current process and future directions. Am J Gastroenterol 2014, 109(5):620–623.

2. Elhassa GO: Drug Development: Stages of Drug Development. Journal of Pharmacovigilance 2015, 03(03).

3. Zurdo J: Developability assessment as an early de-risking tool for biopharmaceutical development. Pharmaceutical Bioprocessing 2013, 1(1):29–50.

4. DiMasi JA, Grabowski HG, Hansen RW: Innovation in the pharmaceutical industry: New estimates of R&D costs. J Health Econ 2016, 47:20–33.

5. DiMasi JA, Hansen RW, Grabowski HG: The price of innovation: new estimates of drug development costs. Journal of Health Economics 2003, 22(2):151–185.

6. Vanhaelen Q: Computational Methods for Drug Repurposing; 2019.

7. Lysenko A, Sharma A, Boroevich KA, Tsunoda T: An integrative machine learning approach for prediction of toxicity-related drug safety. Life Sci Alliance 2018, 1(6):e201800098.

8. Karimi M, Wu D, Wang Z, shen Y: DeepAffinity: Interpretable Deep Learning of Compound Protein Affinity through Unified Recurrent and Convolutional Neural Networks. 2018.

9. Riddick G, Song H, Ahn S, Walling J, Borges-Rivera D, Zhang W, Fine HA: Predicting in vitro drug sensitivity using Random Forests. Bioinformatics 2011, 27(2):220–224.

10. Agarwal S, Dugar D, Sengupta S: Ranking chemical structures for drug discovery: a new machine learning approach. J Chem Inf Model 2010, 50(5):716–731.

11. Napolitano F, Zhao Y, Moreira VM, Tagliaferri R, Kere J, D’Amato M, Greco D: Drug repositioning: a machine-learning approach through data integration. J Cheminform 2013, 5(1):30.

12. Kim E, Choi AS, Nam H: Drug repositioning of herbal compounds via a machine-learning approach. BMC Bioinformatics 2019, 20(Suppl 10):247.

13. Li Z, Wang Y, Xie Y, Zhang L, Dai Z, Zou X: Predicting the binding affinities of compound–protein interactions by random forest using network topology features. Analytical Methods 2018, 10(34):4152–4161.

14. Chen X, Liu M, Gilson MK: BindingDB: a web-accessible molecular recognition database. Comb Chem High Throughput Screen 2001, 4(8):719–725.

15. Chen X, Lin Y, Liu M, Gilson MK: The Binding Database: data management and interface design. Bioinformatics 2002, 18(1):130–139.

16. Gilson MK, Liu T, Baitaluk M, Nicola G, Hwang L, Chong J: BindingDB in 2015: A public database for medicinal chemistry, computational chemistry and systems pharmacology. Nucleic Acids Res 2016, 44(D1):D1045–1053.

17. Liu T, Lin Y, Wen X, Jorissen RN, Gilson MK: BindingDB: a web-accessible database of experimentally determined protein-ligand binding affinities. Nucleic Acids Res 2007, 35(Database issue):D198–201.

18. Chen X, Lin Y, Gilson MK: The binding database: Overview and user’s guide. Biopolymers 2001, 61(2):127–141.

19. Morley-Smith AC, Mills A, Jacobs S, Meyns B, Rega F, Simon AR, Pepper JR, Lyon AR, Thum T: Circulating microRNAs for predicting and monitoring response to mechanical circulatory support from a left ventricular assist device. European journal of heart failure 2014, 16(8):871–879.

20. McGregor MJ, Pallai PV: Clustering of Large Databases of Compounds:⍰ Using the MDL “Keys” as Structural Descriptors. Journal of Chemical Information and Computer Sciences 1997, 37(3):443–448.

21. Durant JL, Leland BA, Henry DR, Nourse JG: Reoptimization of MDL keys for use in drug discovery. J Chem Inf Comput Sci 2002, 42(6):1273–1280.

22. Gautam R, Seider WD: Computation of phase and chemical equilibrium: Part Local and constrained minima in Gibbs free energy. AIChE Journal 1979, 25(6):991–999.

23. Smola AJ, Schölkopf B: A tutorial on support vector regression. Statistics and Computing 2004, 14(3):199–222.

24. Gaulton A, Hersey A, Nowotka M, Bento AP, Chambers J, Mendez D, Mutowo P, Atkinson F, Bellis LJ, Cibrian-Uhalte E et al: The ChEMBL database in 2017. Nucleic Acids Res 2017, 45(D1):D945–D954.

25. Gaulton A, Bellis LJ, Bento AP, Chambers J, Davies M, Hersey A, Light Y, McGlinchey S, Michalovich D, Al-Lazikani B et al: ChEMBL: a large-scale bioactivity database for drug discovery. Nucleic Acids Res 2012, 40(Database issue):D1100–1107.

26. Krasner J: Drug-Protein Interaction. Pediatric Clinics of North America 1972, 19(1):51–63.

27. Akoglu H: User’s guide to correlation coefficients. Turk J Emerg Med 2018, 18(3):91–93.

28. Spearman C: The Proof and Measurement of Association between Two Things. The American Journal of Psychology 1987, 100(3/4):441–471.

29. Hauke J, Kossowski T: Comparison of Values of Pearson’s and Spearman’s Correlation Coefficients on the Same Sets of Data. Quaestiones Geographicae 2011, 30(2):87–93.

30. Fieller EC, Hartley HO, Pearson ES: Tests for Rank Correlation Coefficients. I. Biometrika 1957, 44(3/4):470–481.

31. Erb I, Notredame C: How should we measure proportionality on relative gene expression data? Theory Biosci 2016, 135(1-2):21–36.

32. Lovell D, Pawlowsky-Glahn V, Egozcue JJ, Marguerat S, Bahler J: Proportionality: a valid alternative to correlation for relative data. PLoS Comput Biol 2015, 11(3):e1004075.

33. Quinn TP, Richardson MF, Lovell D, Crowley TM: propr: An R-package for Identifying Proportionally Abundant Features Using Compositional Data Analysis. Sci Rep 2017, 7(1):16252.

34. Girvan M, Newman ME: Community structure in social and biological networks. Proc Natl Acad Sci U S A 2002, 99(12):7821–7826.

35. Silverman LS, Caldwell JP, Greenlee WJ, Kiselgof E, Matasi JJ, Tulshian DB, Arik L, Foster C, Bertorelli R, Monopoli A et al: 3H-[1,2,4]-Triazolo[5,1-i]purin-5-amine derivatives as adenosine A2A antagonists. Bioorg Med Chem Lett 2007, 17(6):1659–1662.

36. Albizu L, Holloway T, Gonzalez-Maeso J, Sealfon SC: Functional crosstalk and heteromerization of serotonin 5-HT2A and dopamine D2 receptors. Neuropharmacology 2011, 61(4):770–777.

37. Wei Y, Pattingre S, Sinha S, Bassik M, Levine B: JNK1-mediated phosphorylation of Bcl-2 regulates starvation-induced autophagy. Mol Cell 2008, 30(6):678–688.

38. Bruey JM, Bruey-Sedano N, Luciano F, Zhai D, Balpai R, Xu C, Kress CL, Bailly-Maitre B, Li X, Osterman A et al: Bcl-2 and Bcl-XL regulate proinflammatory caspase-1 activation by interaction with NALP1. Cell 2007, 129(1):45–56.

39. Wu XY, Zhang CX, Deng LC, Xiao J, Yuan X, Zhang B, Hou ZB, Sheng ZH, Sun L, Jiang QC et al: Overexpressed D2 Dopamine Receptor Inhibits Non-Small Cell Lung Cancer Progression through Inhibiting NF-kappaB Signaling Pathway. Cell Physiol Biochem 2018, 48(6):2258–2272.

40. Wu DM, Wang S, Wen X, Han XR, Wang YJ, Shen M, Fan SH, Zhuang J, Zhang ZF, Shan Q et al: Inhibition of microRNA-200a Upregulates the Expression of Striatal Dopamine Receptor D2 to Repress Apoptosis of Striatum via the cAMP/PKA Signaling Pathway in Rats with Parkinson’s Disease. Cell Physiol Biochem 2018, 51(4):1600–1615.

41. Stam NJ, Van Huizen F, Van Alebeek C, Dijkema R, Tonnaer JADM, Olijve W: Genomic organization, coding sequence and functional expression of human 5-HT2 and 5-HT1A receptor genes. European Journal of Pharmacology: Molecular Pharmacology 1992, 227(2):153–162.

42. Cussac D, Boutet-Robinet E, Ailhaud MC, Newman-Tancredi A, Martel JC, Danty N, Rauly-Lestienne I: Agonist-directed trafficking of signalling at serotonin 5-HT2A, 5-HT2B and 5-HT2C-VSV receptors mediated Gq/11 activation and calcium mobilisation in CHO cells. Eur J Pharmacol 2008, 594(1-3):32–38.

43. Villar VA, Jones JE, Armando I, Palmes-Saloma C, Yu P, Pascua AM, Keever L, Arnaldo FB, Wang Z, Luo Y et al: G protein-coupled receptor kinase 4 (GRK4) regulates the phosphorylation and function of the dopamine D3 receptor. J Biol Chem 2009, 284(32):21425–21434.

44. Patrignani P, Patrono C: Aspirin and Cancer. J Am Coll Cardiol 2016, 68(9):967–976.

45. Warner TD, Mitchell JA: Cyclooxygenase-3 (COX-3): filling in the gaps toward a COX continuum? Proc Natl Acad Sci U S A 2002, 99(21):13371–13373.

46. Seshasai SR, Wijesuriya S, Sivakumaran R, Nethercott S, Erqou S, Sattar N, Ray KK: Effect of aspirin on vascular and nonvascular outcomes: meta-analysis of randomized controlled trials. Arch Intern Med 2012, 172(3):209–216.

47. Penning TM, Burczynski ME, Jez JM, Hung CF, Lin HK, Ma H, Moore M, Palackal N, Ratnam K: Human 3alpha-hydroxysteroid dehydrogenase isoforms (AKR1C1-AKR1C4) of the aldo-keto reductase superfamily: functional plasticity and tissue distribution reveals roles in the inactivation and formation of male and female sex hormones. Biochem J 2000, 351(Pt 1):67–77.

48. Dufort I, Rheault P, Huang XF, Soucy P, Luu-The V: Characteristics of a highly labile human type 5 17beta-hydroxysteroid dehydrogenase. Endocrinology 1999, 140(2):568–574.

49. Ji Q, Aoyama C, Nien YD, Liu PI, Chen PK, Chang L, Stanczyk FZ, Stolz A: Selective loss of AKR1C1 and AKR1C2 in breast cancer and their potential effect on progesterone signaling. Cancer Res 2004, 64(20):7610–7617.

50. Ji Q, Chang L, VanDenBerg D, Stanczyk FZ, Stolz A: Selective reduction of AKR1C2 in prostate cancer and its role in DHT metabolism. Prostate 2003, 54(4):275–289.

51. Tai H-L, Lin T-S, Huang H-H, Lin T-Y, Chou M-C, Chiou S-H, Chow K-C: Overexpression of aldo-keto reductase 1C2 as a high-risk factor in bladder cancer. Oncology Reports 2007.

52. Byrns MC, Penning TM: Type 5 17beta-hydroxysteroid dehydrogenase/prostaglandin F synthase (AKR1C3): role in breast cancer and inhibition by non-steroidal anti-inflammatory drug analogs. Chem Biol Interact 2009, 178(1-3):221–227.

53. Ertl P, Rohde B, Selzer P: Fast calculation of molecular polar surface area as a sum of fragment-based contributions and its application to the prediction of drug transport properties. J Med Chem 2000, 43(20):3714–3717.

54. Pajouhesh H, Lenz GR: Medicinal chemical properties of successful central nervous system drugs. NeuroRx 2005, 2(4):541–553.

55. Stefanis L: alpha-Synuclein in Parkinson’s disease. Cold Spring Harb Perspect Med 2012, 2(2):a009399.

56. Caillet-Boudin ML, Buee L, Sergeant N, Lefebvre B: Regulation of human MAPT gene expression. Mol Neurodegener 2015, 10:28.

57. Weingarten MD, Lockwood AH, Hwo SY, Kirschner MW: A protein factor essential for microtubule assembly. Proc Natl Acad Sci U S A 1975, 72(5):1858–1862.

58. Lockhart A: Imaging Alzheimer’s disease pathology: one target, many ligands. Drug Discov Today 2006, 11(23-24):1093–1099.

59. Ye L, Velasco A, Fraser G, Beach TG, Sue L, Osredkar T, Libri V, Spillantini MG, Goedert M, Lockhart A: In vitro high affinity alpha-synuclein binding sites for the amyloid imaging agent PIB are not matched by binding to Lewy bodies in postmortem human brain. J Neurochem 2008, 105(4):1428–1437.

