## Supplementary Information for "Prediction of drug-protein interaction and drug repositioning using machine learning model"

### Supporting Information

#### SI Methods

#### Calculation of protein similarities

To discover the potential target for drug repositioning, we examined the protein similarity by calculation of the predicted $-logK_i$ values of drugs between proteins. The predicted $-logK_i$ values were processed before calculation of similarities because most of them were relatively small and could not be informative. Since we wanted to focus on the comparisons of those drugs with higher values, the abundance of these noises may interfere with the correlations of the higher predictions between proteins. Hence, we calculated the standard scores (*z*-score) of predicted $-logK_i$ for each protein model and the z-scores with values larger than 2 were retained, while the others were assigned values of 0.

Here we introduced proportionality $\rho$ [1-3] to indicate the correlations between proteins and built a proportionality-based protein-protein network. Given a matrix of *D* features measured across *N* samples, before calculating the correlations between samples, the features of each sample are computed by a centered log-ratio transformation (clr), which scales each subject vector by its geometric mean (g(x)). For features $[x_1, \dots , x_i, \dots, x_D]$ in each sample, the transformation is defined as

$$\mathrm{clr}(x) = \left[\ln\frac{x_1}{\mathrm{g}(x)}, \dots, \ln\frac{x_i}{\mathrm{g}(x)}, \dots, \ln\frac{x_D}{\mathrm{g}(x)}\right].$$

The log-ration transformations to each sample results in a new matrix $A$ with $N$ samples and $D$ features. Then the proportionalities $\rho$ between two samples $A_i$ and $A_j$ are calculated basing on log-ratio variance according to

$$\rho_p(A_i, A_j) = 1 - \frac{var(A_i - A_j)}{var(A_i) + var(A_j)},$$

which ranges from -1 (perfect reciprocality) to +1 (perfect proportionality).

Since the value of proportionality ranges from -1 to 1, which was not appropriate when being used as a weight to build a network, we rescaled the range from 0 to 1 by multiplying the value by 0.5 and then add 0.5 to it. To note, the rescaled values with smaller than 0.5 represented negative correlation, greater than 0.5 positive correlation, and 0.5 uncorrelated.

**GO term similarity**

To see whether two proteins have a correlation in their roles in biological processes, we used their annotations from Gene Ontology (GO) (Dataset S3) to calculate the similarity between them. The similarity between proteins $A$ and $B$ was calculated as the Jaccard similarity coefficient and was defined as the size of the intersection of GO terms of both proteins divided by the size of the union of GO terms of both proteins

$$J(A, B) = \frac{|A \cap B|}{|A \cup B|}.$$

Since the ontologies of GO are structured as a hierarchical graph, relations between terms should be taken into account when calculating similarities. For each biological process term of a protein, all of its ancestors were considered as the terms of the protein to eliminate the calculation bias caused by hierarchy.

When evaluating the indirect associations between proteins through the shortest paths in the eBC network, we had to take edge number and the medium proteins into account to calculate the GO term similarities. The Jaccard similarity coefficient that may include more than two members was calculated as

$$J(A,X) = \frac{|A \cap B \cap \ldots \cap X|}{|(A \cap B) \cup (B \cap C) \cup \ldots \cup (W \cap X)|},$$

where $A$ and $X$ were the starting and ending proteins respectively, and $[B, C, \ldots, W]$ were the proteins that the shortest path past.

### SI Figures

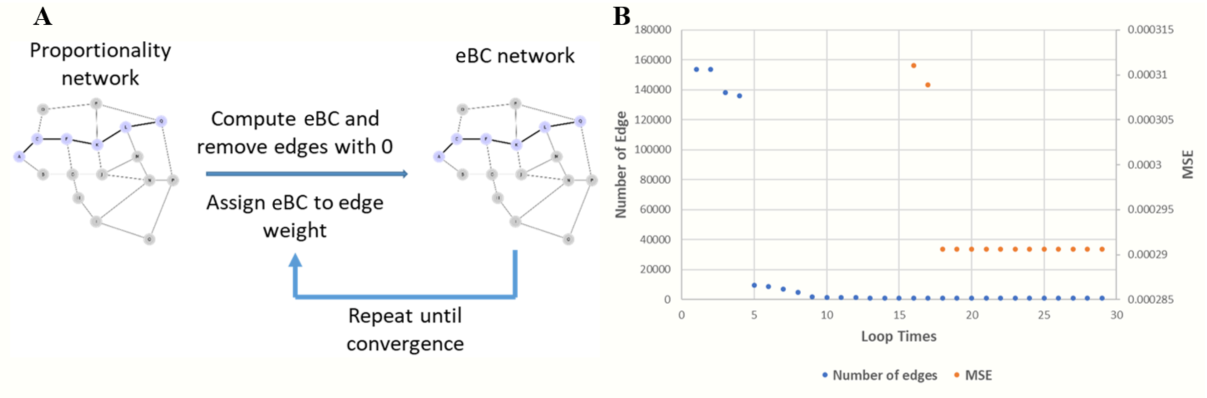


**Figure S1. Construction of eBC-based protein-protein network.**

(A) The edges with eBC values of 0 were removed and the eBC values were used as the new weights until the network converged. (B) Record of edge numbers and MSE in eBC cycles. The edge number and the MSE value stayed stable since the 10th and 18th cycle respectively

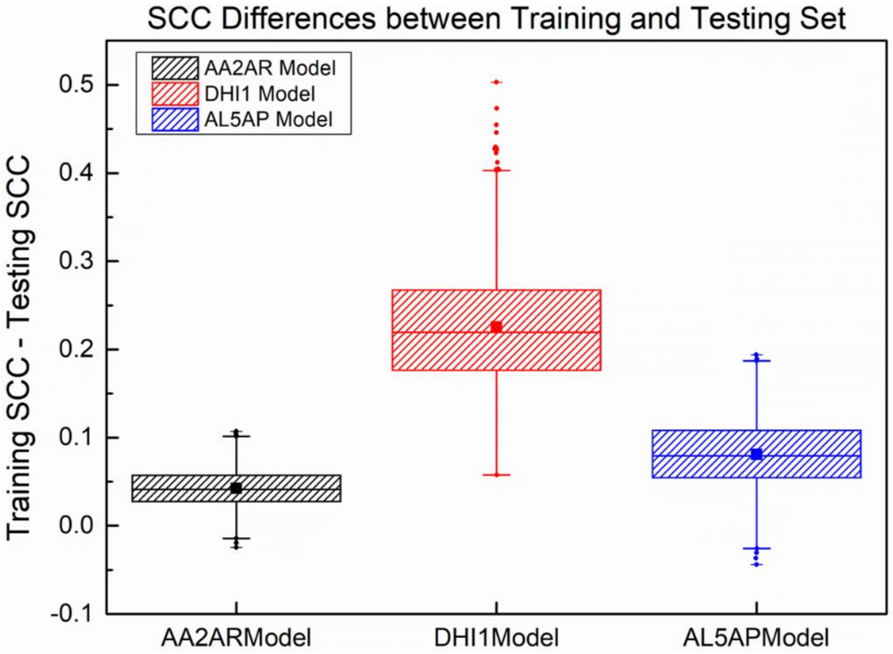


**Figure S2. Distributions of 1,000 SCC differences between training and testing set**

The AA2AR model had the most drugs data among the three models and had the smallest difference between training and testing SCC, while the DHI1 model had the least drugs and the largest difference.

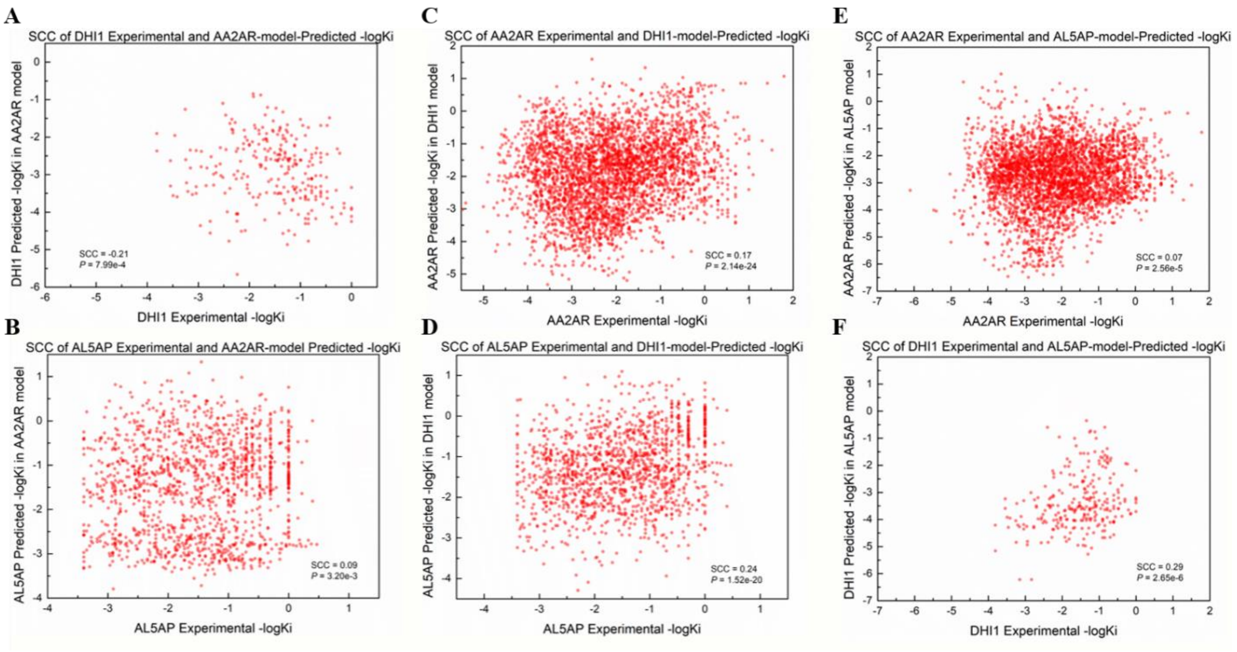


**Figure S3. Model specificities Evaluation**

(A) (B) AA2AR model predicting DHIA and AL5AP drugs. (C) (D) DHIA model predicting AA2AR and AL5AP drugs. (E) (F) AL5AP model predicting AA2AR and DHIA drugs. Binding predictions using models of other proteins showed low correlations with the experimental data.

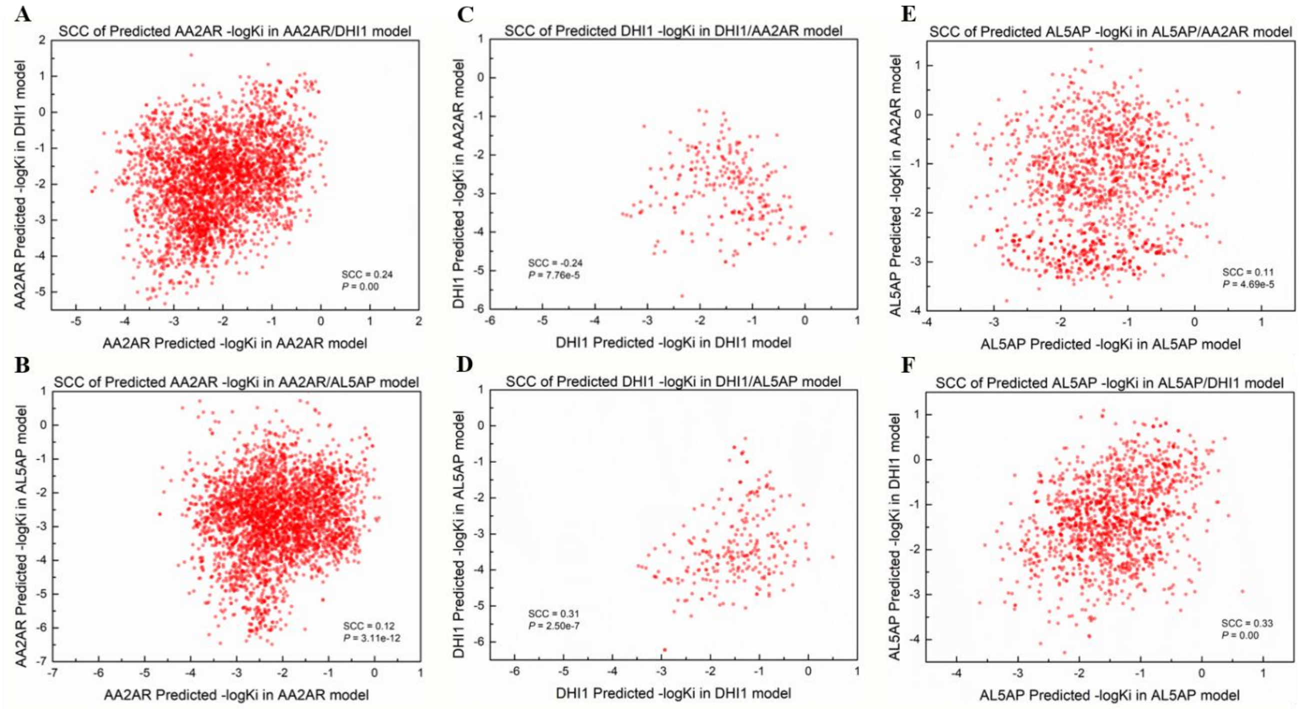


**Figure S4. Models specificities evaluation**

(A) AA2AR drugs predicted in AA2AR and DHI1 models. (B) AA2AR drugs predicted in AA2AR and AL5AP models. (C) DHI1 drugs predicted in DHI1 and AA2AR models. (D) DHI1 drugs predicted in DHI1 and AL5AP models (E) AL5AP drugs predicted in AL5AP and AA2AR models. (F) AL5AP drugs predicted in AL5AP and DHI1 models. Binding predictions using their own models showed low correlation with those using models of other proteins.

### SI Tables

**Table S1. Physicochemical properties used in this study [4, 5]**

| Feature | Description | Feature | Description |
|---|---|---|---|
| Full_mwt | Molecular weight | Mw_freebase | Molecular weight of the parent form of a molecule[I]. |
| Mw_monoisotopic | The sum of the masses of the most abundant isotopes in the compound | AlogP | Partition coefficient, is used to evaluate the lipophilicity or hydrophobic of a molecule[II]. |
| Acd_logP | The octanol/water partition coefficient using a fragment-based method developed by ACDlabs. | Acd_logD | The calculated octanol/water Distribution Coefficient at pH7.4. |
| Aromatic_rings | The number of aromatic rings atoms in a molecule. | Heavy_atoms | The number of non-hydrogen atoms in a molecule. |
| Hba | The number of hydrogen bond acceptors. | Hbd | The number of hydrogen bond donors. |
| Hba_lipinski | The count of nitrogen and oxygen atoms in the molecule. | Hbd_lipinski | The count of hydrogens attached to nitrogen or oxygen atoms. |
| Num_ro5_violations | The number of properties in a molecule that fails to meet the definition in Lipinski's rule of 5 (RO5). | Num_lipinski_ro5_violations | Replacing "hba" and "hbd" by "hba_lipinski" and "hbd_lipinski" in "num_ro5_violations". |
| Rtb | The number of rotatable bonds in a molecule. | PSA | The polar surface area[III]. |
| Qed_weighted | A quantitative estimate of drug-likeness[IV]. | | |

[I]A parent structure is a compound that consists of an unbranched chain of skeletal atoms, according to IUPAC nomenclature. In pharmacokinetics, logP is a significant indicator of ADME properties, since a drug needs to be lipophilic to pass through lipid bilayers in the intestinal epithelium after taken orally, but still hydrophilic enough to partition out again. As for pharmacodynamics, more lipophilicity means more toxic because a drug with high logP tends to retained longer within the body.

[II]AlogP is measured as the ratio of concentrations of molecular species in different aqueous media, usually octanol and water. The larger the logP value is, the higher the affinity of a molecule to the organic phase is.

[III]PSA is defined as the sum of surface area over all polar atoms, chiefly oxygen and nitrogen and their attached hydrogen atoms.

[IV]The value range from 0-1 where 1 is the most drug-like and 0 the least drug-like[6].

### SI Datasets

**Dataset S1. The binding database**

**Dataset S2. The ChEMBL database**

**Dataset S3. The gene ontology resource**

### References


1. Erb I, Notredame C: **How should we measure proportionality on relative gene expression data?** *Theory Biosci* 2016, **135**(1-2):21-36.
2. Lovell D, Pawlowsky-Glahn V, Egozcue JJ, Marguerat S, Bahler J: **Proportionality: a valid alternative to correlation for relative data**. *PLoS Comput Biol* 2015, **11**(3):e1004075.
3. Quinn TP, Richardson MF, Lovell D, Crowley TM: **propr: An R-package for Identifying Proportionally Abundant Features Using Compositional Data Analysis**. *Sci Rep* 2017, **7**(1):16252.
4. Gaulton A, Hersey A, Nowotka M, Bento AP, Chambers J, Mendez D, Mutowo P, Atkinson F, Bellis LJ, Cibrian-Uhalte E *et al*: **The ChEMBL database in 2017**. *Nucleic Acids Res* 2017, **45**(D1):D945-D954.
5. Gaulton A, Bellis LJ, Bento AP, Chambers J, Davies M, Hersey A, Light Y, McGlinchey S, Michalovich D, Al-Lazikani B *et al*: **ChEMBL: a large-scale bioactivity database for drug discovery**. *Nucleic Acids Res* 2012, **40**(Database issue):D1100-1107.
6. Bickerton GR, Paolini GV, Besnard J, Muresan S, Hopkins AL: **Quantifying the chemical beauty of drugs**. *Nat Chem* 2012, **4**(2):90-98.
